# Global vertebrate hotspots

**DOI:** 10.64898/2026.05.13.724836

**Authors:** Harith Farooq, Michael B. J. Harfoot, Carsten Rahbek, Piero Visconti, Jonas Geldmann

## Abstract

Effective biodiversity conservation requires tools that can identify priority areas under growing human pressures. Building on the concept of global biodiversity hotspots, we present a transparent and repeatable approach to mapping conservation priorities using data for 33,604 species of terrestrial vertebrates from the IUCN Red List. This framework expands the taxonomic scope of previous efforts and integrates updated information on key human-driven threats to biodiversity. We identify that around 13% of Earth’s terrestrial surface qualifies as vertebrate conservation hotspots, often shaped by distinct combinations of species groups and threats. These results highlight the need for tailored, context-specific conservation strategies. By providing a robust method to guide spatial prioritization, our work supports more effective implementation of conservation targets in a rapidly changing world.

## Introduction

Biodiversity is declining at an alarming rate (Cardinale et al., 2012; Ceballos & Ehrlich, 2002; Pimm et al., 2014) with as much as 30% of assessed species being threatened with extinction (IPBES, 2019). This decline have intensified over the last decades and is linked to human activities (Díaz et al., 2019). The loss of species and ecosystems not only represent an impoverishment of nature, but also, in many places, jeopardizes our own livelihoods (Rockstrom et al., 2009). A key step to safeguard biodiversity is identifying those areas of most importance to biodiversity and ensuring their persistence in perpetuity through protected and conserved areas or other conservation actions (Margules & Pressey, 2000; Rodrigues & Cazalis, 2020). Global approaches to identify areas of conservation importance, while rarely suitable for supporting local scale decision-making, can help guide such processes by synthesizing important knowledge and providing large-scale perspectives (Chaplin-Kramer et al., 2022). One hugely influential approach to identify areas of priority for biodiversity conservation has been the identification of global ‘biodiversity hotspots’ (Fjeldsaå et al., 1997; Myers, 1988; Myers et al., 2000). These hotspots combine two fundamental aspects related to priority setting: 1) they have high level of endemism, making them irreplaceable for the conservation of several species and 2) they have experienced large historic losses of natural vegetation (Myers et al., 2000).

The Biodiversity hotspots concept has been instrumental in shaping global conservation priorities and help identify some of the areas of the world most in need of conservation attention (Dalton, 2000; Sloan et al., 2014). However, these ‘hotspots’ have also been critiqued for relying too heavily on poor data, and a simplistic approach to map broad regions of biodiversity importance (Jepson & Canney, 2001; Kareiva & Marvier, 2003; Marchese, 2015; Norman, 2003). In particular, the original biodiversity hotspots ignored all threat other than extent of native vegetation being lost and biodiversity importance was based only on the amount of endemic plants (Myers et al., 2000), with no strict methodology or threshold to identify areas with high endemism. This resulted in maps with inadequate taxonomical scope (i.e. only plants), course resolution, and based on methods with limited transparency and repeatability. In addition, since the first publication of ‘biodiversity hotspots’, over two decades ago, nature conservation (Mace, 2014) and conservation planning (Adams et al., 2004; Iwamura et al., 2010; Naidoo et al., 2006; Wilson et al., 2006) has changed significantly. This change has in particular resulted in a much greater understanding of the role of people as potential ‘victims’ or ‘winners’ of conservation interventions (Brockington & Wilkie, 2015; Naidoo et al., 2019). Thus, conservation strategies cannot only be evaluated on their ability to safeguard biodiversity but also need to incorporate both the potential costs and benefits that these strategies might incur on those people and communities living in and around the areas set aside for biodiversity (Carpenter et al., 2009; Ghoddousi et al., 2021).

Here we develop a framework for locating ‘biodiversity hotspots’ building on the original concept of identifying areas that are of importance for biodiversity and under high threat. Towards this, we draw one of the most important knowledge products in conservation (Brooks et al., 2015; Juffe-Bignoli et al., 2016), using updated maps of all terrestrial amphibians, birds, mammals, and reptiles (n = 33,604) extracted from the IUCN Red List of threatened species (BirdLife International and NatureServe, 2022; International Union for Conservation of Nature, 2022). Our approach combines two specific spatial products – threats measured as the probability of a random species in a given location being affected by a specific threat using the method developed by (Harfoot et al., 2021) and biodiversity importance measured through WEGE which combines the rarity and extinction risk of species in a given location (Farooq et al., 2020). To map threats, we use the main direct drivers of biodiversity loss globally (IPBES, 2019): agriculture, logging, urbanization, over-exploitation and hunting, climate change, invasive species, and pollution (Farooq et al., 2024; Harfoot et al., 2021). While we acknowledge that our approach is limited by only including vertebrates, it addresses some of the most important criticism related to the original ‘biodiversity hotspots’, including their low resolution, limited taxonomic scope, and lack of reproducibility. More importantly, our framework can be adapted to other resolutions and to also account for other taxa and threats, as new data become available. The vertebrate hotspots may potentially guide national governments, national and international NGOs and other stakeholders in identifying regions that require further conservation attention; including monitoring, financing, law enforcement, and possibly protected and conserved areas.

## Results

We identified ca. 13% of Earth’s terrestrial surface (18.7 million km^2^) as vertebrate hotspots (Fig. 1). These are primarily located in the tropics and dominated by broadleaf forests and mangroves with between 34.2% and 53% of their total area identified as a hotspot respectively. While the vertebrate hotspots align reasonably well with the biodiversity hotspots (64% overlap compared the Null expectation of 19% if hotspots were located randomly), ca, 1/3 of the vertebrate hotspots fall outside the original ones with some important deviations, in particular in the Caucasus and South American dry grasslands (Fig. 1). Conversely, Madagascar, the tropical forests in Africa and Southeast Asia, New Zealand, Central America, and the Polynesian islands had a disproportionately high coverage of vertebrate hotspots (Fig. 1).

**Figure 1.**
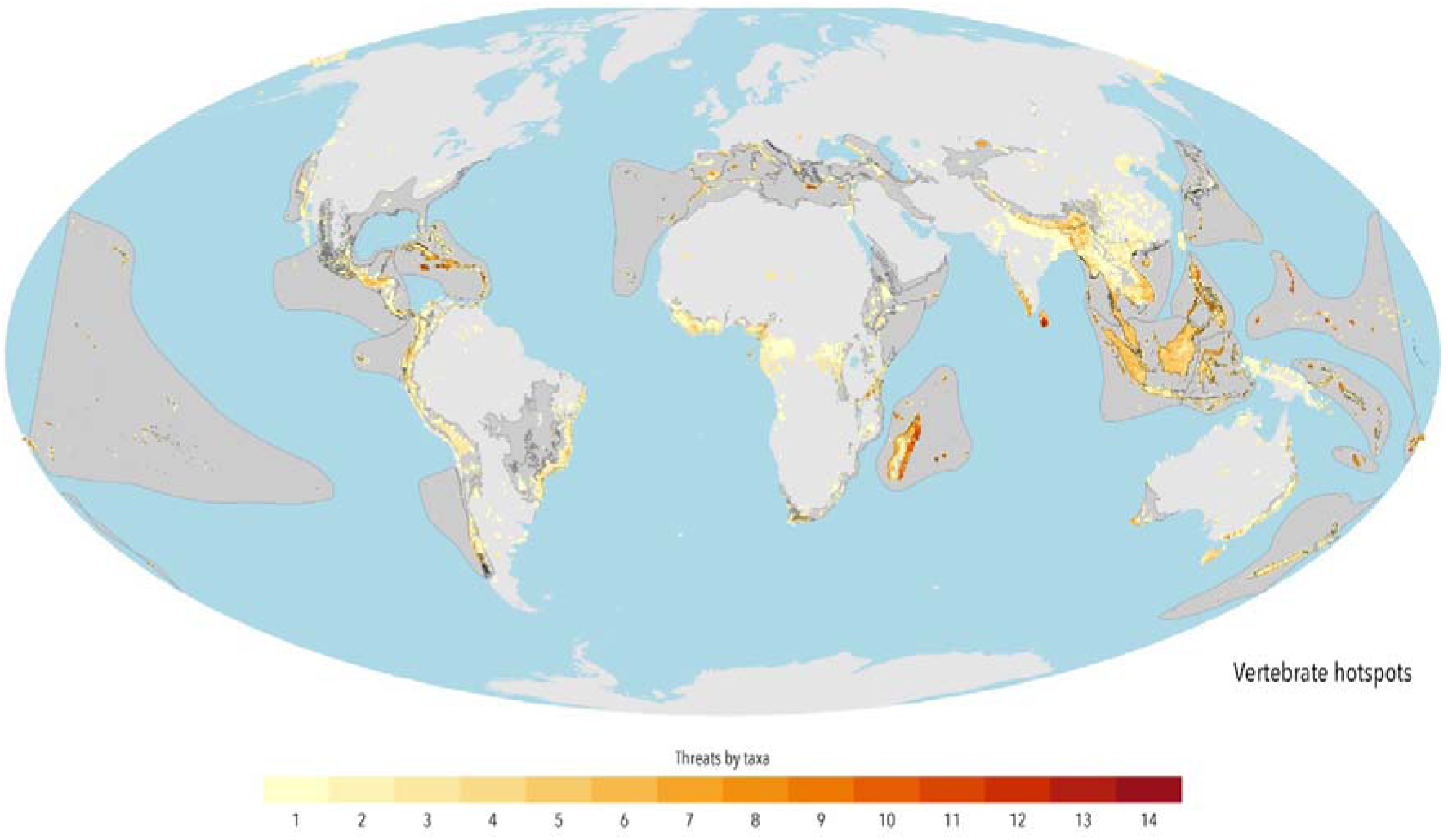
Map of vertebrate hotspots. The distribution of grid cells that match our thresholds to be considered a vertebrate hotspot, coloured by the cumulative number of threats by taxa, where 1 is the minimum possible value and 28 is the maximum possible value, which would have been translated into the four taxonomic groups threatened by all the seven analyzed threats. The grid cells were placed on a map of the world’s land in light grey and in darker grey the 36 known biodiversity hotspots.

To control for global latitudinal gradient of species richness, which can skew priorities towards the tropics (Rohde, 1992), we also identify vertebrate hotspots within each Wallace’s Zoogeographical regions (Holt et al., 2013). Using this approach, we highlight areas in Central and East Asia as well as much of both the European and North African Mediterranean coast, which is not identified using our global approach, but which emerges as potential vertebrate hotspots at the regional level (Fig. 2). Mammals (47%) and reptiles (39%) triggered the most hotspots grids, followed by birds (38%) and amphibians (33%) (Fig. 2).

**Figure 2.**
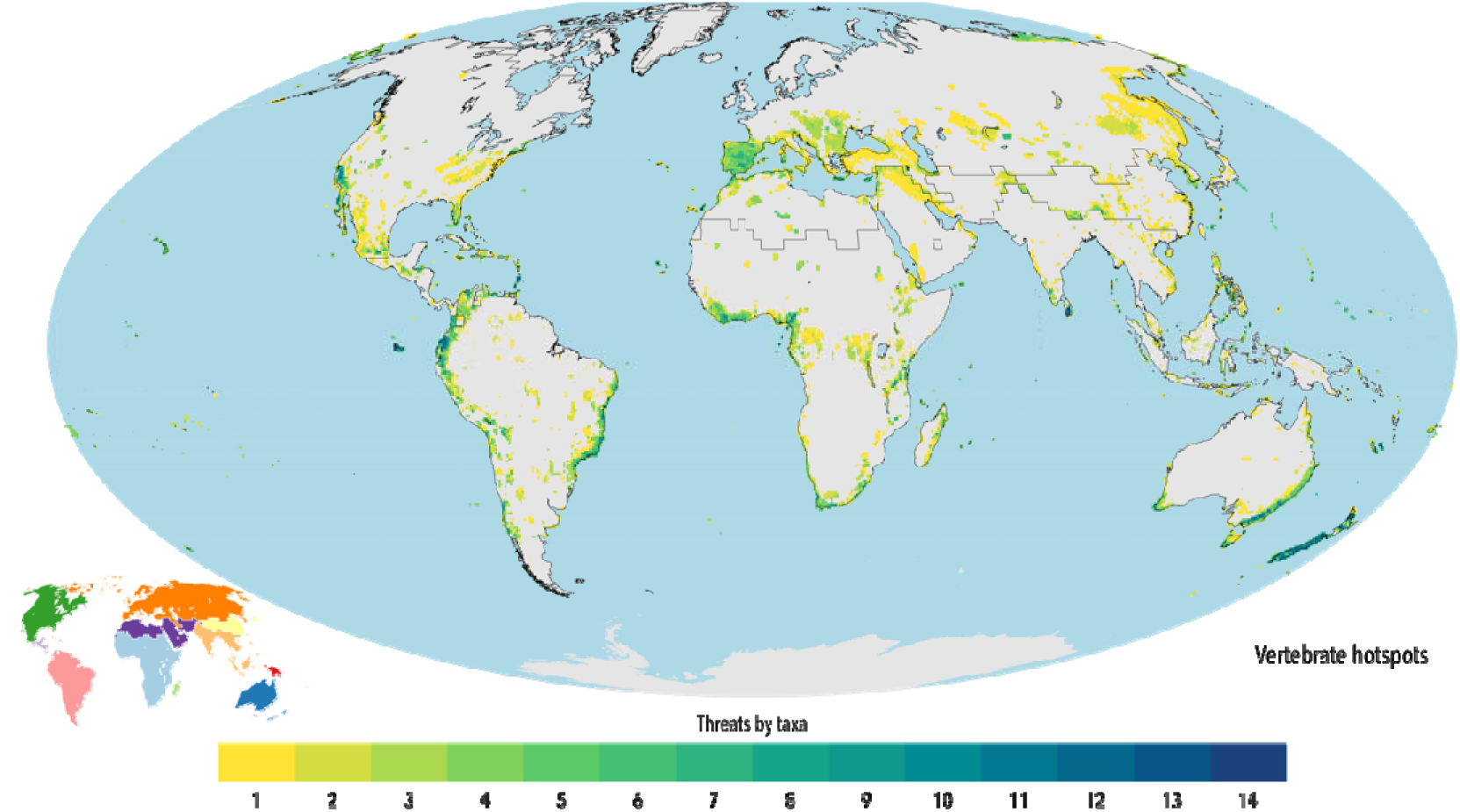
Distribution of vertebrate hotspots calculated for each Wallace’s Zoogeographic Region separately. The map shows the top 10 decile WEGE cells that overlap with at the 10 decile likelihood of threat for at least one threat but calculated separately for each of the 11 realms illustrated in the lower left corner. The grid cells were coloured based on the cumulative number of threats by taxa.

**Figure 3.**
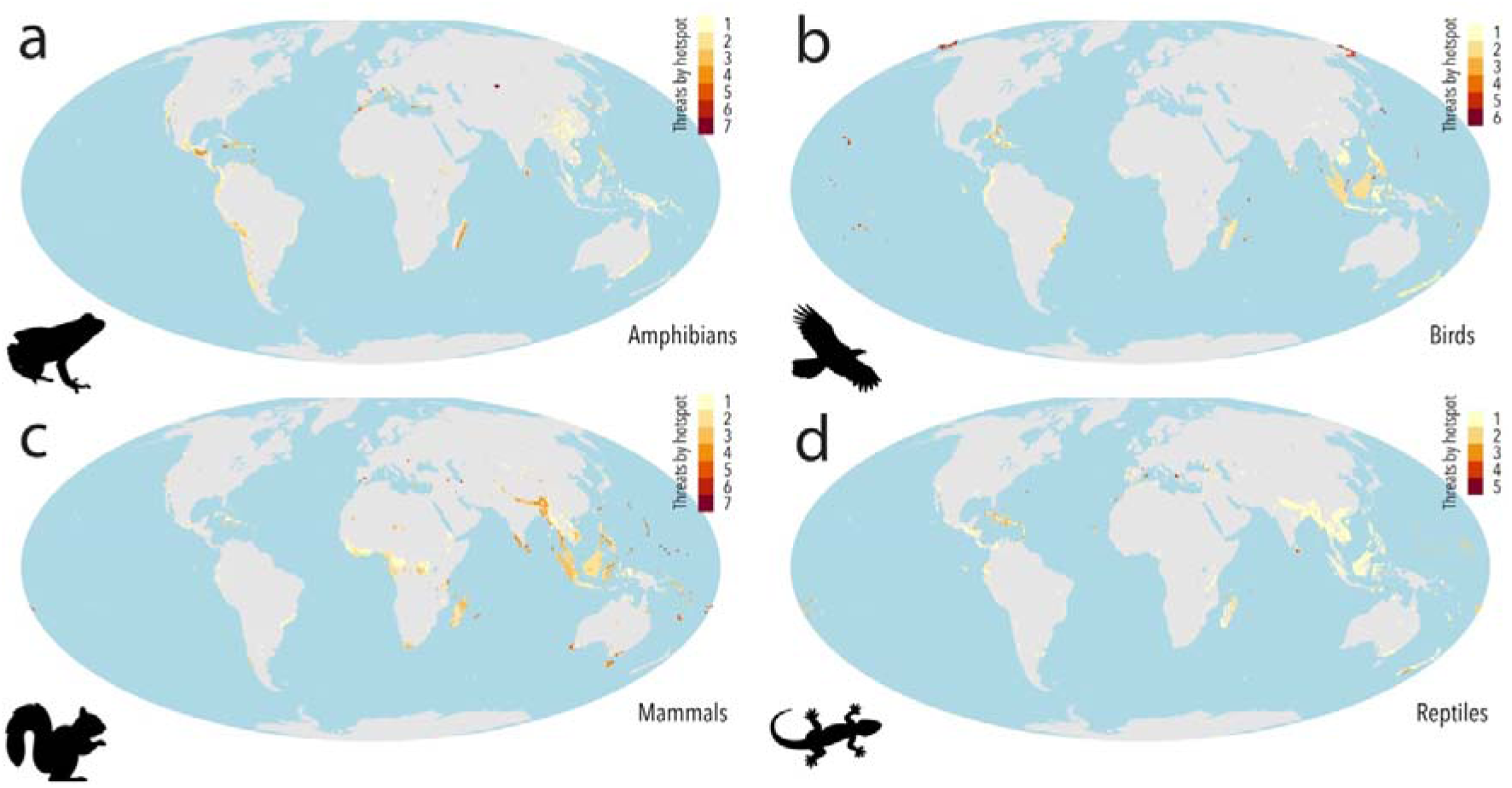
The distribution of grid cells that match our thresholds to be considered a vertebrate hotspot, coloured by the cumulative number of threats for each of amphibians, birds, mammals and reptiles respectively **a-d**. Here 1 is the minimum possible value and 7 is the maximum possible value.

Habitat destruction and degradation was the most significant threat, with logging (a trigger in 66% of the hotspots – 15,425,000 km^2^) as the primary driver, followed by agriculture (a trigger in 44% of the hotspots – 11,267,500 km^2^), and urbanization (a trigger in 40% of the hotspots – 5,345,000 km^2^; Fig. 4a, b, g). The methodological approach behind the vertebrate hotspots also reveal that other threats not included in the original biodiversity hotspots are both important and pervasive. Invasive species (a trigger in 30% of the hotspots – 5,367,500 km^2^), hunting, (a trigger in 17% of the hotspots – 3,187,500 km^2^), climate change (a trigger in 16% of the hotspots – 3,325,000 km^2^), and finally pollution (a trigger in 10% of the hotspots – 1,842,500 km^2^) were, thus, all major threats in those areas most important for biodiversity (Fig. 4c-f). Besides logging, which was the most important threat for all four taxa, amphibians and reptiles were particularly affected by invasive species, while mammals were particularly affected by agriculture, urbanization, and hunting (Fig. 4g).

**Figure 4.**
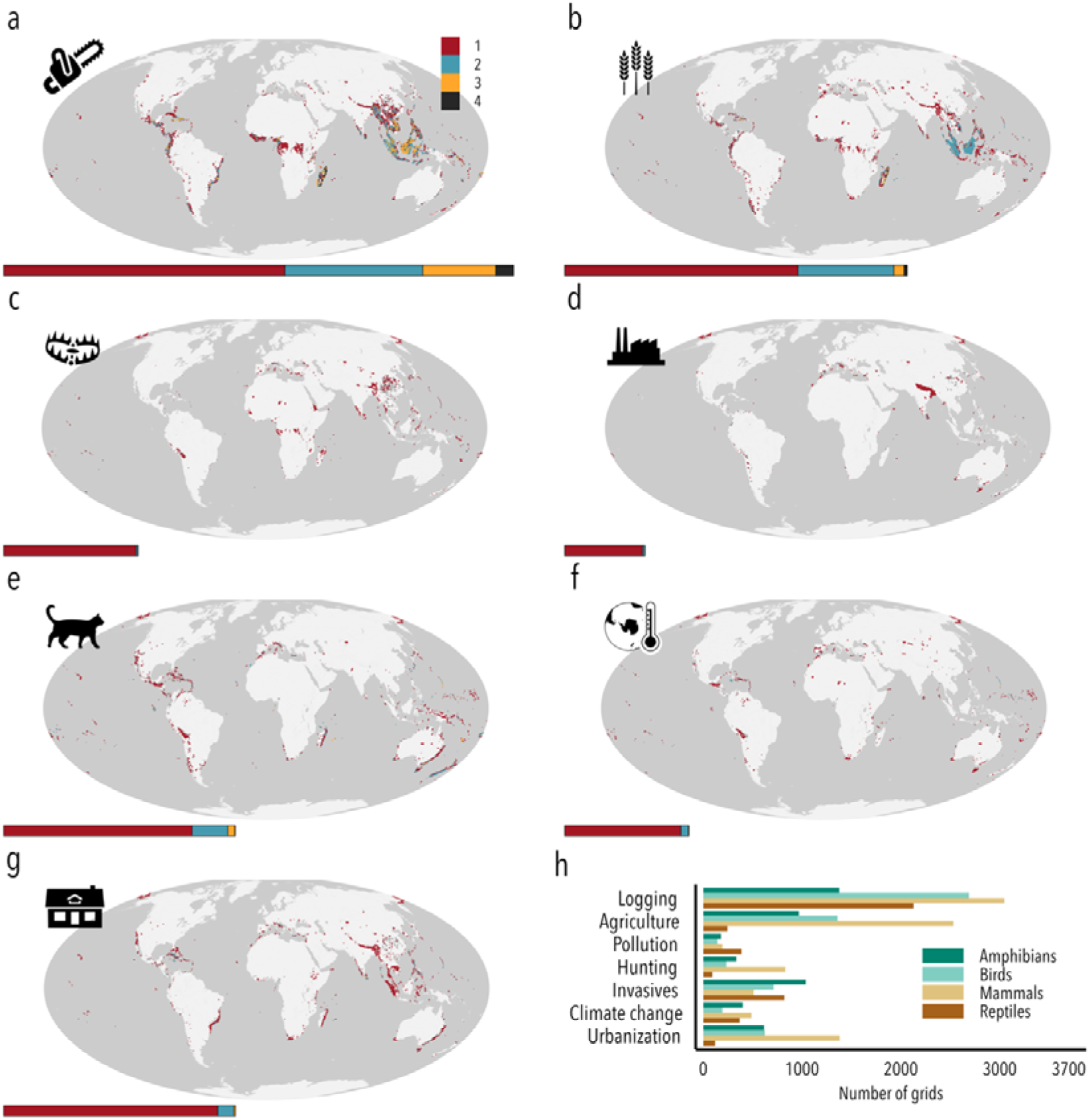
Maps a-g showing the top 10 deciles for each threat that overlap with the top 10 WEGE cells. All maps are coloured by the number of taxonomic groups triggering each cell. In dark grey are cells that are in the top 10 decile of threat that overlap with the top 10 WEGE cells – these areas fit the biological requirements to become a hotspot but are yet not under the top 10 decile of threat. Under each map there is a stacked bar showing the cumulative number of grid cells triggered by each threat in relation to logging. In h is the number of hotspot grids for each threat broken down by taxonomic group. Threats affect taxonomic groups differently, e.g. hunting is a more important threat of mammals when compared to reptiles, while climate change affects amphibians more than birds.

While our vertebrate hotspots only account for 13% of surface of Earth, we found that over a third of the global human population live inside of them (Fig. 5). This pattern is particularly pronounced in the tropics, with for example the hotspots occurring in tropical and subtropical moist broadleaf forests covering only 0.6% (1 million km^2^) of Earth’s terrestrial surface but being home to approximately 23% of the human population (Fig. 5).

**Figure 5.**
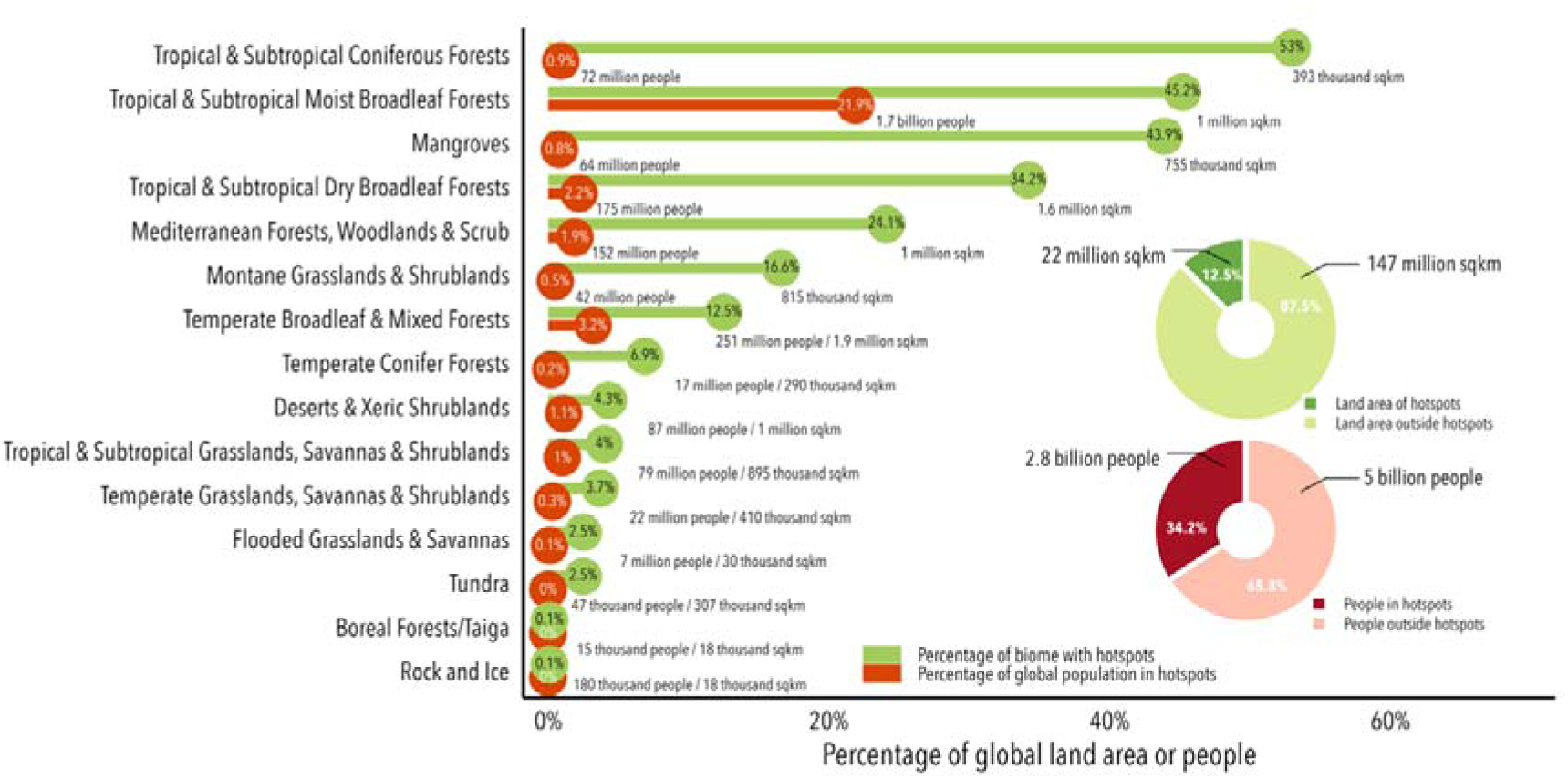
The percentage of global area and global population contained in the vertebrate hotspots, in each biome. Over a third of the global population live within vertebrate hotspots and approximately a fourth are in hotspots situated in the Tropical and Subtropical Moist Broadleaf Forests biome.

## Discussion

The vertebrate hotspots provide a timely update and addition to one of the most important global priority layers for biodiversity conservation. We show that while there are areas across the world that are hotspots for multiple taxa and threats, this is not the case in the majority of areas. Indeed, areas of importance to biodiversity are often affected by a complex combination of threats targeting specific taxa. The lack of overlap between threats and taxa highlights the strength of mapping specific threats rather than relying on measures that combine the impact of multiple drivers into one compound metric (Allan et al., 2019). While protected and conserved areas might be useful tools against land use changes and some illegal harvesting (Geldmann et al., 2013; Graham et al., 2021), they might be less ideal for addressing more pervasive threats such as pollution or climate change (Mu et al., 2021). Similarly, invasive species eradication, while effective in particular on islands (Langhammer et al., 2024), might require resources that are likely not realistic to allocate at a more continental scale (McConnachie et al., 2012). Thus, conservation strategies instead of extrapolating from specific threats or taxa to formulate global strategies, they must build on more holistic approaches, that incorporates the complexity of how different taxa responds to different threats as well as the integration of local perspectives and solutions (Bowler et al.; Farooq et al., 2024; Hof et al., 2011).

### A methodological improvement

The framework presented here for locating ‘biodiversity hotspots’ builds on the original concept of identifying areas that are of importance for biodiversity and under high threat. Importantly, our proposed methodology is pixel-based, rather than relying on courser biogeographical regions. This approach allows the mapping of hotspots at a finer resolution and their update as new and more accurate data becomes available, thereby ensuring that repeatability and transparency is incorporated in how these hotspots are identified. Since the original biodiversity hotspots were published, decades have passed, our knowledge has improved, and new threats have emerged and intensified (Richardson et al., 2023). Regardless of the method used to map hotspots, places of high endemism but low threat two decades ago, may have become current hotspots due to cumulative and accelerating pressures. Therefore, mapping biodiversity hotspots as a continuously updated process, helps prevent hotspots from becoming outdated. Our approach also expands the taxonomic coverage, while also addressing some of the criticisms that have been raised against the original biodiversity hotspots. Despite the vertebrate hotspots aligning reasonably well with the biodiversity hotspots, approximately a third of the vertebrate hotspots fall outside the biodiversity ones. This discrepancy might be driven by these habitats importance for plants (Noss, 2012; Wilson et al., 2012), which was the primary biodiversity component of the biodiversity hotspots (Myers et al., 2000), but omitted here due to the lack of comprehensive data. The vertebrate hotspots reveal that other threats not included in the original biodiversity hotspots, such as invasive species, hunting, climate change, and pollution, are both important and widespread. Together, these threats mean that a percentage of 19% of the identified vertebrate hotspots would be omitted if habitat loss was to be used as the sole proxy for threats to biodiversity. Besides logging, which was the most important threat for all four taxa, amphibians and reptiles were particularly affected by invasive species, consistent to previous findings (Holmes et al., 2019; Spatz et al., 2017), while mammals were particularly affected by agriculture, urbanization, and hunting (Tilman et al., 2017). Noteworthy, since over a third of the global human population live in the vertebrate hotspots, we also integrate the potential implications for people by translating our hotspots into priority setting.

### Vertebrate hotspots and people

Although vertebrate hotspots account for just 13% of Earth’s surface, over a third of the global human population lives within these areas. Therefore, the development of conservation strategies that rely on these maps must prioritize the rights and well-being of affected people and communities. Protected areas can benefit peoples and communities (Campos-Silva et al., 2021; den Braber et al., 2018; Li et al., 2024; Naidoo et al., 2019; Oldekop et al., 2016) however, poorly executed conservation measures can also cause harm. Such negative outcomes include the eviction of communities (Lele et al., 2010), the erosion of their rights (Rights and Resources Initiative, 2015) and livelihoods (Brockington & Wilkie, 2015), and not properly integrating indigenous people and local communities in governance and management (McCarthy et al., 2018). Without addressing such factors, any strategy that seeks to expand the footprint of conservation, risks not only having adverse effects on people but also reducing support for, and effectiveness of conservation efforts (Geldmann, 2024). The vertebrate hotspots in several places extend across national borders emphasizing a need for transboundary conservation initiatives (Liu et al., 2020) and a global approach that shares the burdens equitably is needed to secure the most important areas for biodiversity (Shen et al., 2023).

### Limitations

The vertebrate hotspots can serve as a useful component for conservation prioritization and planning through highlighting areas of potential global conservation importance. However, the approach implemented in this study identifies areas at a ∼50x50 km, and thus, is unable to identify specific sites, but rather highlight broader areas that would require more careful assessments based on on-the-ground data and be integrated into a wider plan for protection and conservation action (Palacio et al., 2023). In addition, the focus on both biodiversity importance and threats, means that hotspots type priority maps ignores potential ‘cold spots’ and wilderness area, that harbours important biodiversity but is not currently under threat (Kareiva & Marvier, 2003; Marchese, 2015; Watson et al., 2016), as well as areas of potential importance for nature’s contribution to people (Jung et al., 2021). The vertebrate hotspots presented here rely solely on terrestrial vertebrates which constitute less than 1% of all species (Mora et al., 2011). As a result, they fail to represent the full scope of biodiversity. The exclusion of plants, fungi and invertebrates is likely to skew the selection of sites (Di Marco et al., 2017; Heilmann-Clausen et al., 2015) and their inclusion would likely reveal areas of importance not captured by our maps. Furthermore, while the core data for our analysis is one of the most trusted and tested datasets in conservation (Brooks et al., 2015; Butchart et al., 2005) it is not without its flaws and biases (Cazalis et al., 2022; Rondinini et al., 2014). Of special relevance is the under-representation of some of the most important, but under-sampled, areas for conservation like the tropics (Hughes et al., 2021). Nevertheless, despite these biases, due to the nature of the framework presented here, which species and regions are included when mapping hotspots could be tailored to fit the purpose of each study.

### Final considerations

With the global community committed to ensuring that the integrity, connectivity and resilience of all ecosystems are maintained and that human induced extinction of known threatened species is halted by 2050 (Convention on Biological Diversity, 2022), understanding how to best reduce the loss of areas of high biodiversity importance, including ecosystems of high ecological integrity is vital. By broadening the concept of biodiversity hotspots beyond plants to include vertebrate hotspots, we provide an essential contribution to global conservation prioritization, supporting both international and local planning processes.

However, identifying the right places to conserve is only part of the solution. Equally important is ensuring that conservation actions taken in these sites can reverse current negative biodiversity trends and support the livelihood of people. By integrating vertebrate hotspots into conservation strategies and focusing on their effectiveness, we can make significant strides toward preserving biodiversity and preventing further species decline.

## Methods

To map the threats to the terrestrial vertebrates we downloaded all the range maps of amphibians, birds, mammals, and reptiles from IUCN (International Union for Conservation of Nature, 2022) and BirdLife International and NatureServe (BirdLife International and NatureServe, 2022) and used the IUCN Threats Classification Scheme (Version 3.2), which contains 11 primary threat classes (Salafsky et al., 2008). In total, after removing marine reptiles (i.e. all sea turtles and sea snakes) and all categories other than DD, LC, NT, VU, EN, CR, we were left with 33,604 species (7,209 amphibians, 10,984 birds, 5,608 mammals, and 9,803 reptiles). To limit the effects of commission errors we further limited the ranges to presence for all four taxonomic groups, bird ranges included both presence and breeding ranges.

We then used the distribution ranges and threat information to build threat layers at a resolution of 50 x 50 km for agriculture, climate change, hunting, invasive species, logging, pollution, and urbanization. The probability of threat was calculated as in (Harfoot et al., 2021), using a binomial distribution where the response variable consisted of whether a species were coded as affected by a specific threat or not and weighted by the inverse of the cube root of range size.

To calculate conservation importance of each 50 x 50 km cell, we calculated WEGE scores, using the following formula as in Farooq et al. (2020):

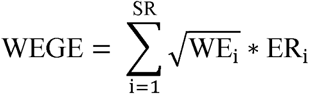

The WEGE value for each cell represents the sum of the square root of the partial weighted endemism value (WEi) multiplied by the probability of extinction (ERi) for each species recorded within a given 50 x 50 km cell. Further details of this calculation are outlined in Farooq et al. (2020). Weighted endemism is inversely proportional to the number of grid cells in which a species occurs within. For extinction risk, the IUCN50 transformation from Davis et al. (2018) was used with the following probabilities: LC = 0.0009, NT = 0.0071, VU = 0.0513, EN = 0.4276, and CR = 0.9688. For DD species the probability of VU was used (0.0513) following Bland et al. (2015) who found that DD species are often assessed as threatened when data for their assessment becomes available. To diminish the effect of species assessed as a single point, all species that had ranges smaller than 20 km^2^ were set to this value.

We then set a threshold of for the top 10 decile of the WEGE metric for each of four taxonomic groups as well as for each of the seven threats as our measure of high biodiversity importance and high threat respectively. Thus, if a cell was both in the top 10 decile for WEGE for at least one taxonomic group and also on the top 10 decile of one of the threats for the respective taxonomic group, the cell would be considered a vertebrate hotspot cell. We generated global maps summing the number of threats by taxa where the maximum value possible to obtain is 28, corresponding to the product of the number of taxonomic groups (4) by the number of threats (7). We repeated our approach of overlap between the top 10 decile of threat and WEGE separately for each of the 11 Wallace’s Zoogeographic Regions obtained in Holt et al. (2013). To quantify the contribution of different threats in general and for each of the taxonomic groups, we counted the number of hotspot cells due to a specific threat and for how many taxa such grid cell was triggered. To compare across taxonomic groups and quantify how each taxonomic group is affected by each threat, we counted the number of hotspots grid cells triggered by each threat.

To calculate the human population counts living inside our hotspot grids, we aggregated the population count (CIESIN, 2018) for the year 2020 from a resolution of ∼1km to a ∼50km to match our grid of hotspots. We then calculated the percentage of the global population and area within the hotspot grids in each of the global biomes obtained in (Dinerstein et al., 2017) (Fig 5). All analysis were conducted in a Mollweide projection using R (R version 4.2.2). The conversion of shapefiles into grids for both species ranges and regions used a touch algorithm. This was used to avoid omission of narrow-ranged species and regions. To analyse the congruence of our hotspots and the current biodiversity hotspots we calculated the random overlap of terrestrial cells with the existing hotspots over 1000 iterations – null hypothesis, and then compared it to our obtained overlap.

In an analysis presented in the supplementary material, we repeated the same steps using ecoregions obtained in (Dinerstein et al., 2017). To do this, we calculated the percentage of ecoregions that have biodiversity hotspot cells, we converted their shape files into 50 km x 50 km grids. Since the ecoregions’ land map omits several islands, we lumped the cells that lack any ecoregion if they were at most 250 km away from the ecoregion. We then calculated the proportion of cells of each ecoregion that have biodiversity hotspot grids.

An important feature of the vertebrate hotspots is that it identifies the specific taxonomic group and threats that trigger each location. Due to this methodological improvement on the biodiversity hotspots, we are able to quantify the differences across the four taxa in terms of what areas were identified as hotspots, with 61% of locations triggered by only one taxon (Fig. 3; Fig. 4). This specificity reinforces that relying on just one species group, risks overlooking important areas for others (Larsen et al., 2012) and also clearly show that our vertebrate hotspots do not replace the original ones based on plant data, but rather compliment them.

## Acknowledgements

H.F. and J.G. were supported by The Danish Independent Research council (grant no. 0165-00018B) and H.F. and C.R. were supported by the Villum Center for Global Mountain Biodiversity.

## Data availability statement

All the data used in this study was obtained from the IUCN Red List of Threatened of Threatened Species (IUCN, 2024) and is freely available for academic research.

## Code availability statement

The code used in this study can be obtained from https://github.com/harithmorgadinho/hotspots.git

## Author contribution statement

J.G. conceived the study; H.F. and J.G designed and implemented the analysis with input from M.B.J.H, C.R. and P.V.; H.F. collected, collated, and organized all data; J.G. and H.F. prepared the manuscript with input from M.B.J.H, C.R. and P.V.

