## Supplementary material for "Global vertebrate hotspots"

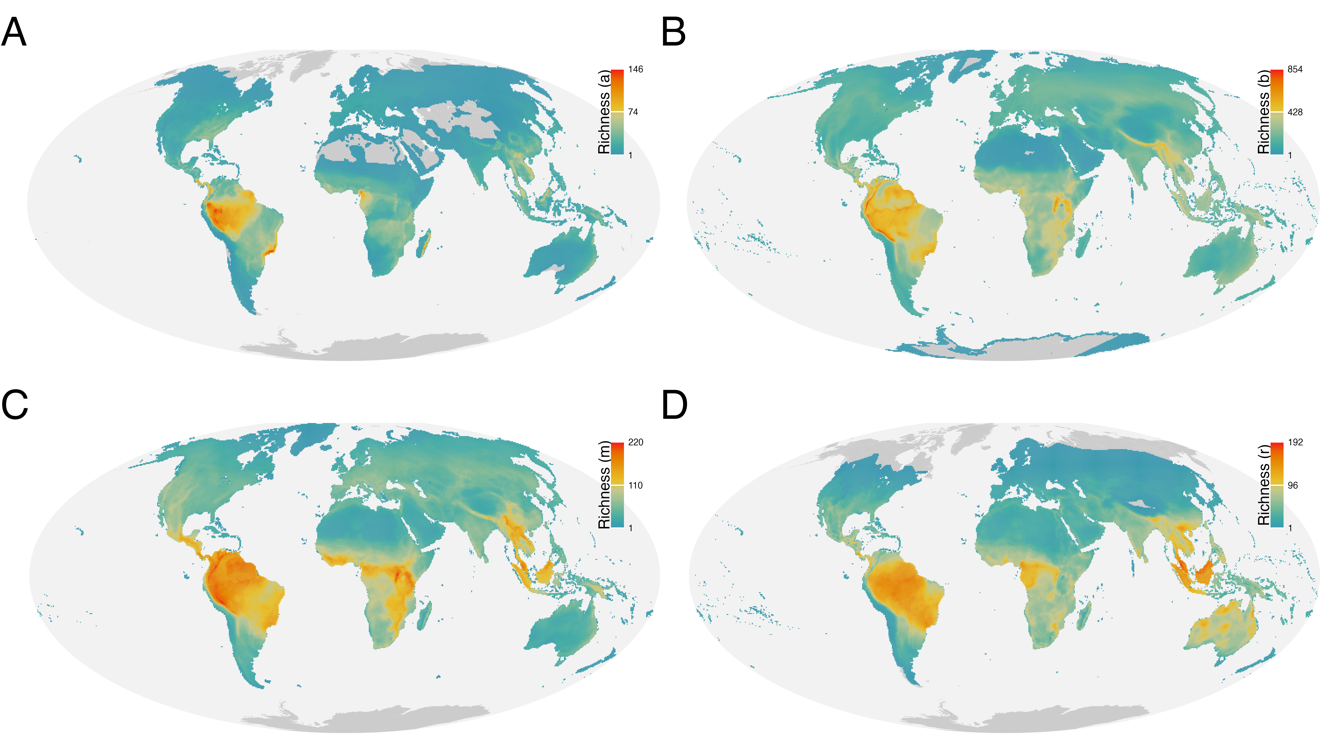

**Figure S1** | Global richness maps of the four taxonomic groups used in the analysis. A. Amphibians, B. Birds, C. Mammals and D. Birds.

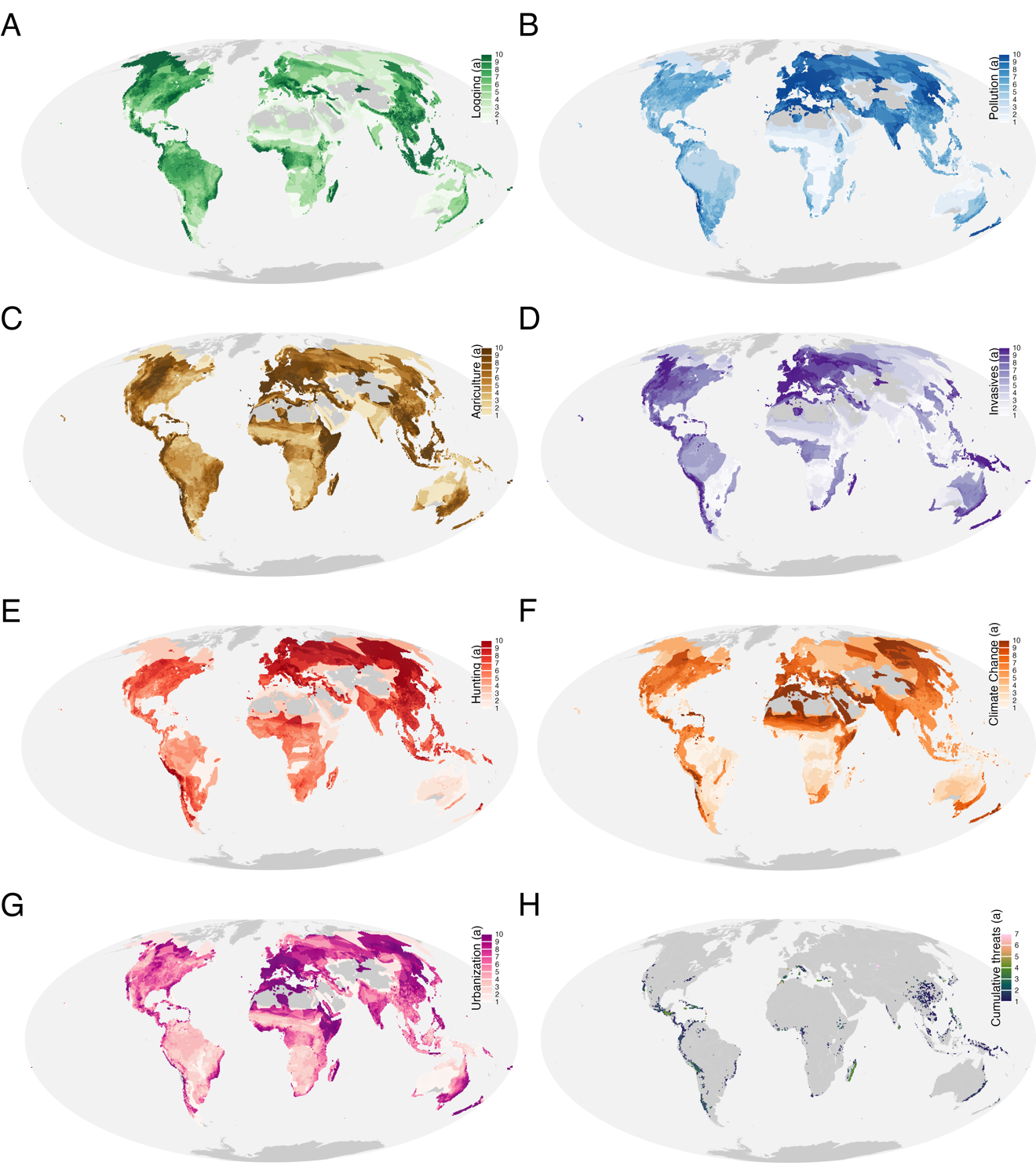

**Figure S2** | Threat maps for amphibians and coloured by deciles. A. Logging, B. Pollution. C. Agriculture, D. Invasive species, E. Hunting, F. Climate change, G. Urbanization and H. The sum of threats for each the value is in the 10th quantile.

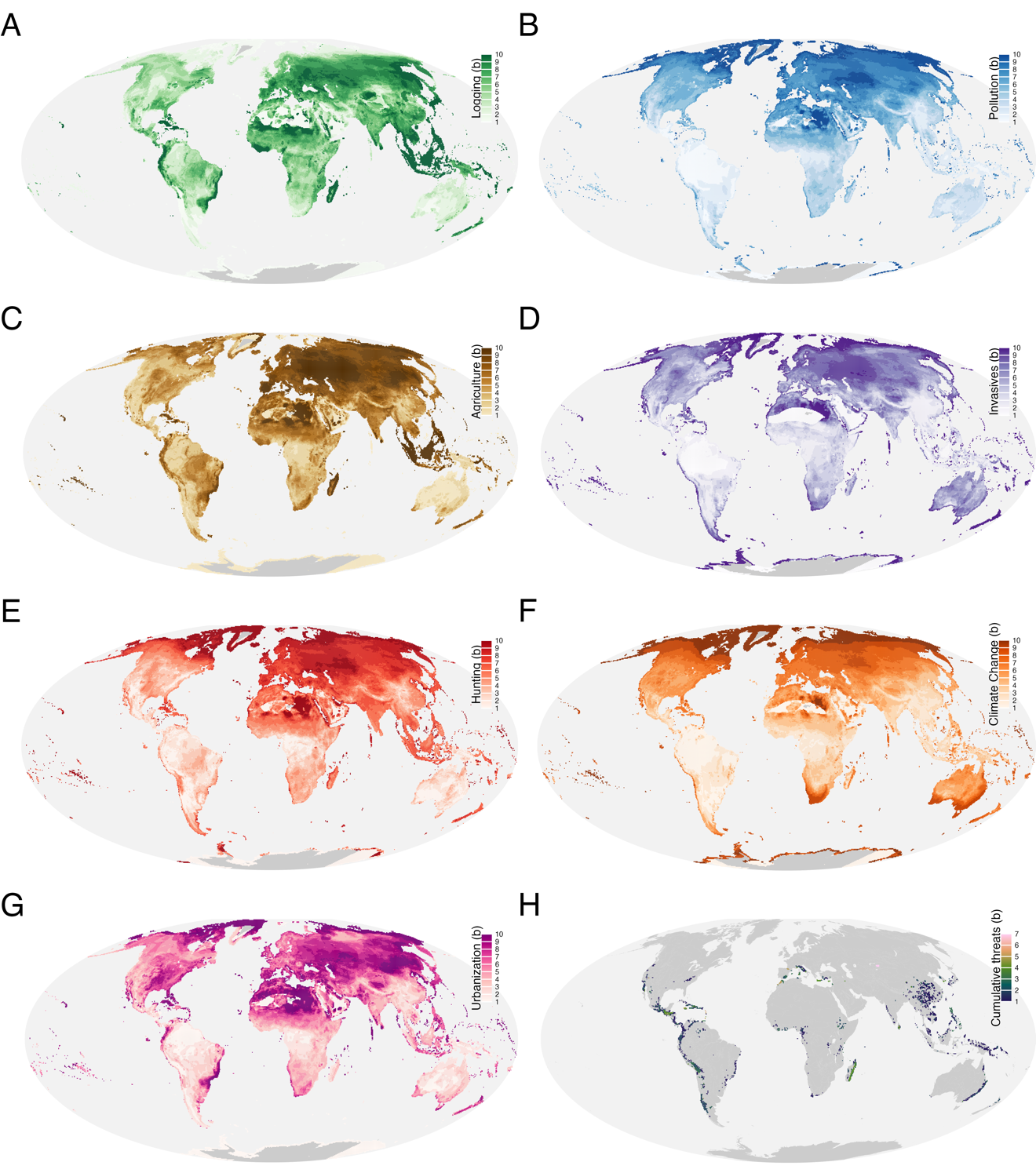

**Figure S3** | Threat maps for birds and coloured by deciles. A. Logging, B. Pollution. C. Agriculture, D. Invasive species, E. Hunting, F. Climate change, G. Urbanization and H. The sum of threats for each the value is in the 10th quantile.

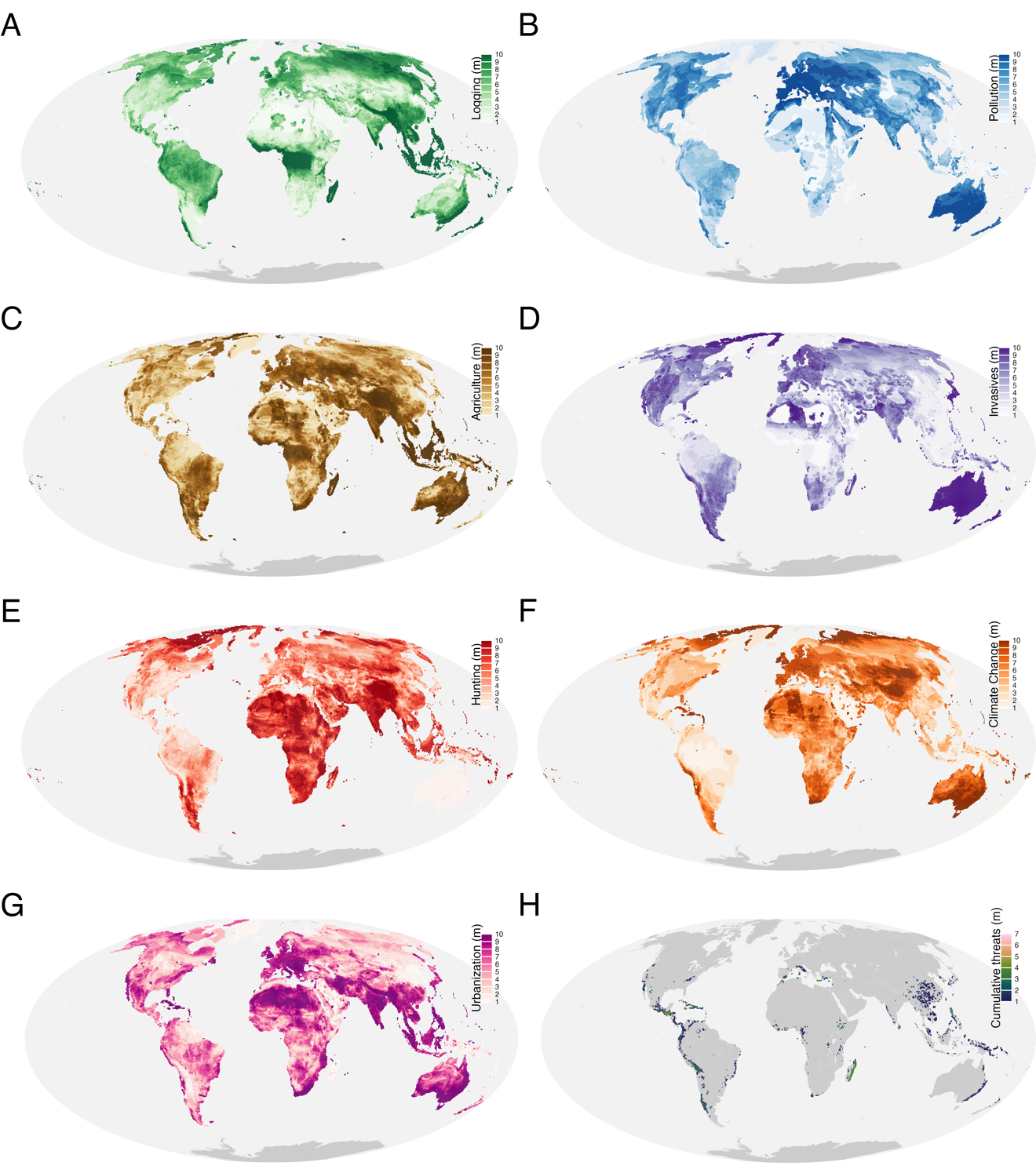

**Figure S4** | Threat maps for mammals and coloured by deciles. A. Logging, B. Pollution. C. Agriculture, D. Invasive species, E. Hunting, F. Climate change, G. Urbanization and H. The sum of threats for each the value is in the 10th quantile.

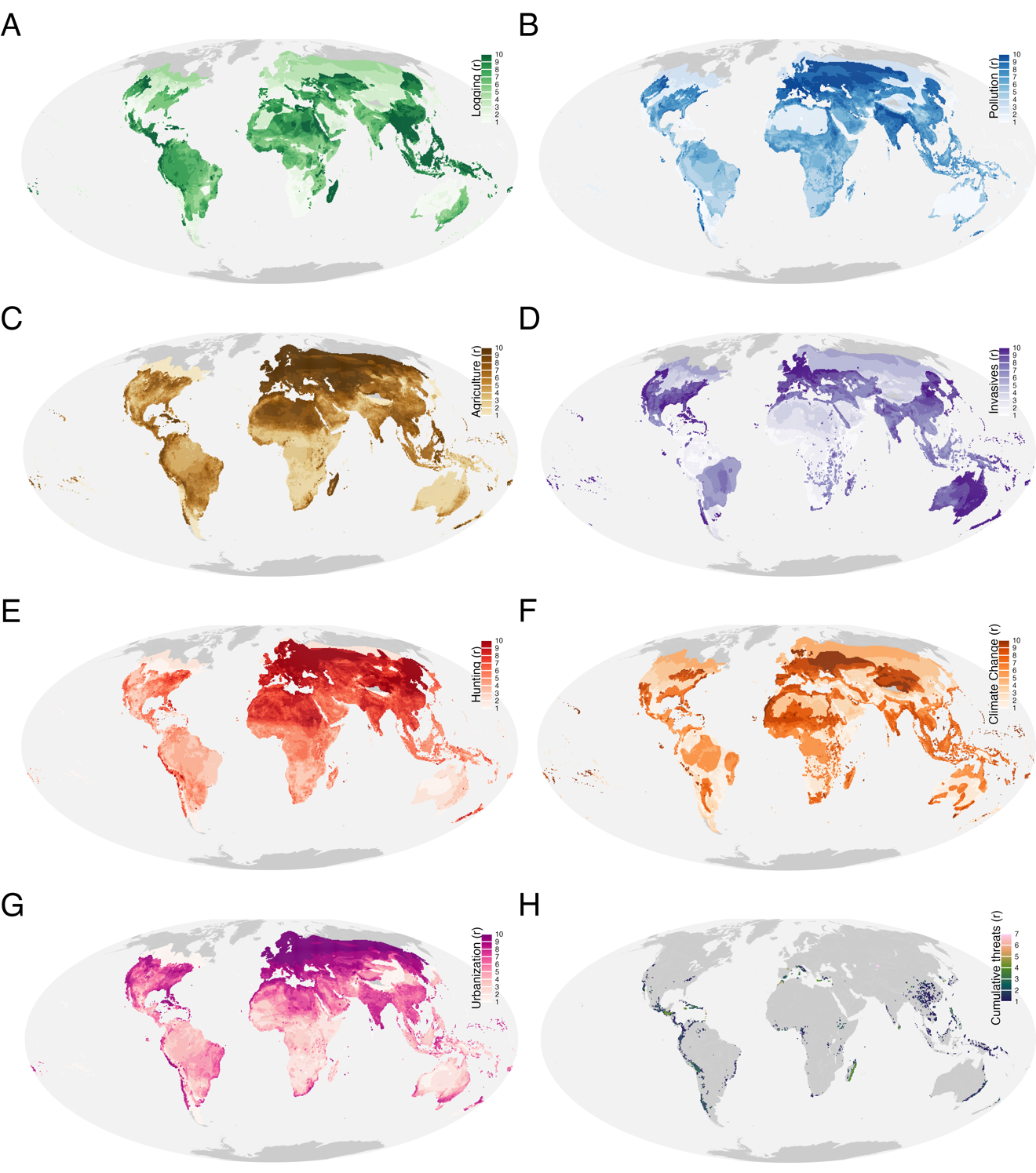

**Figure S5** | Threat maps for reptiles and coloured by deciles. A. Logging, B. Pollution. C. Agriculture, D. Invasive species, E. Hunting, F. Climate change, G. Urbanization and H. The sum of threats for each the value is in the 10th quantile.

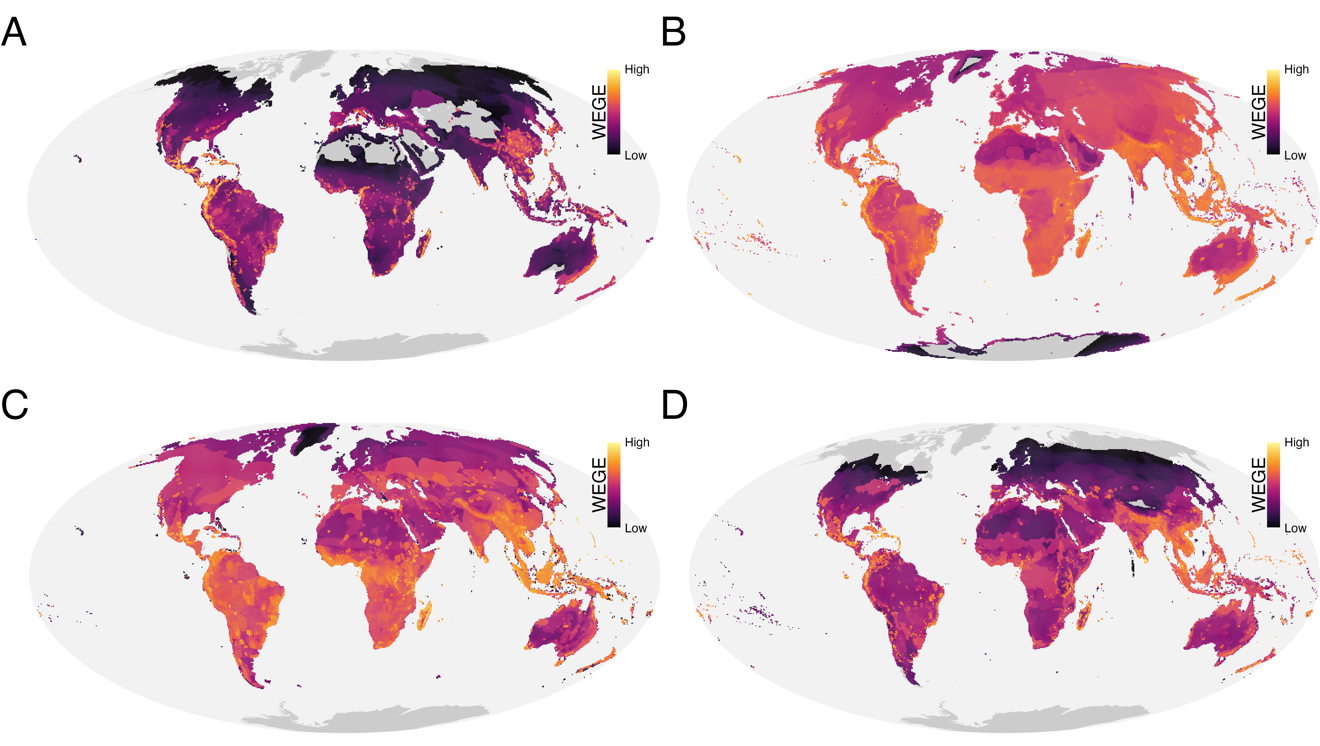

**Figure S6** | Global distribution of WEGE scores (log10 transformed) for A. Amphibians, B. Birds, C. Mammals, D. Reptiles.

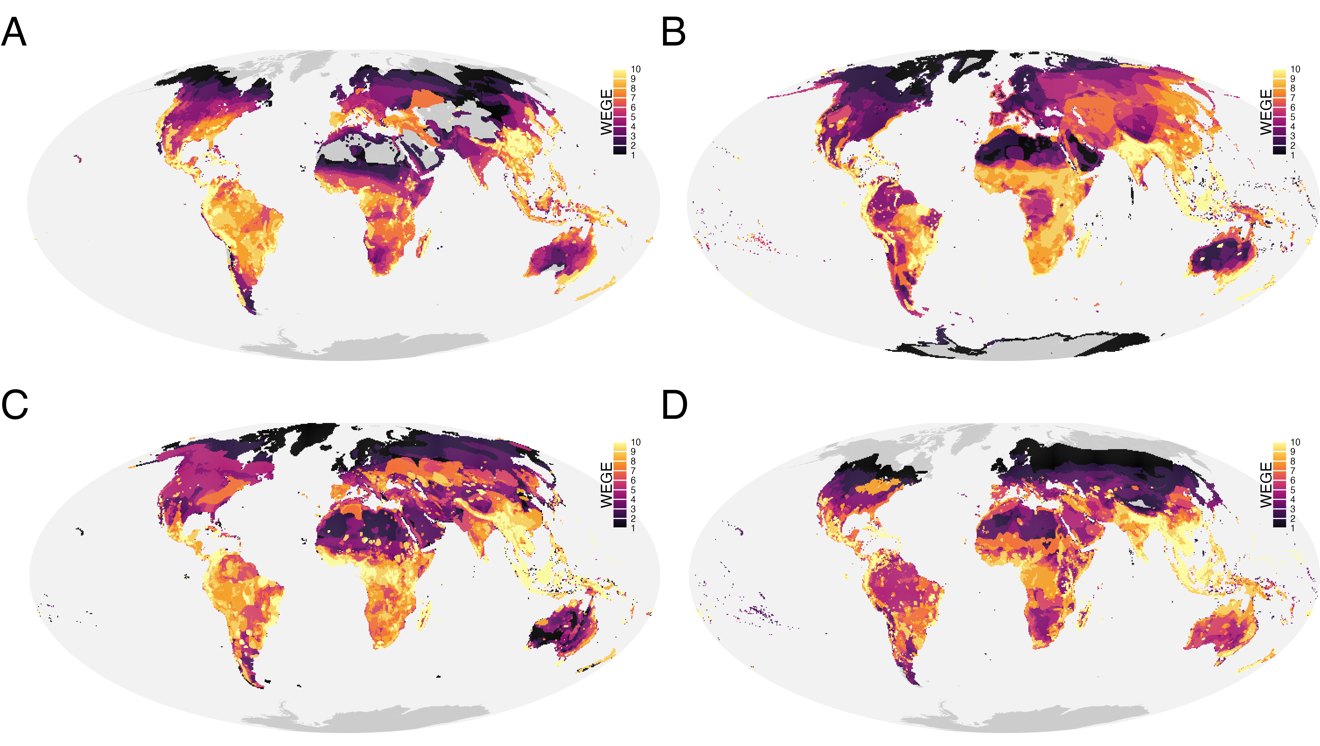

**Figure S7** | Global distribution of WEGE scores coloured by decile for A. Amphibians, B. Birds, C. Mammals, D. Reptiles.

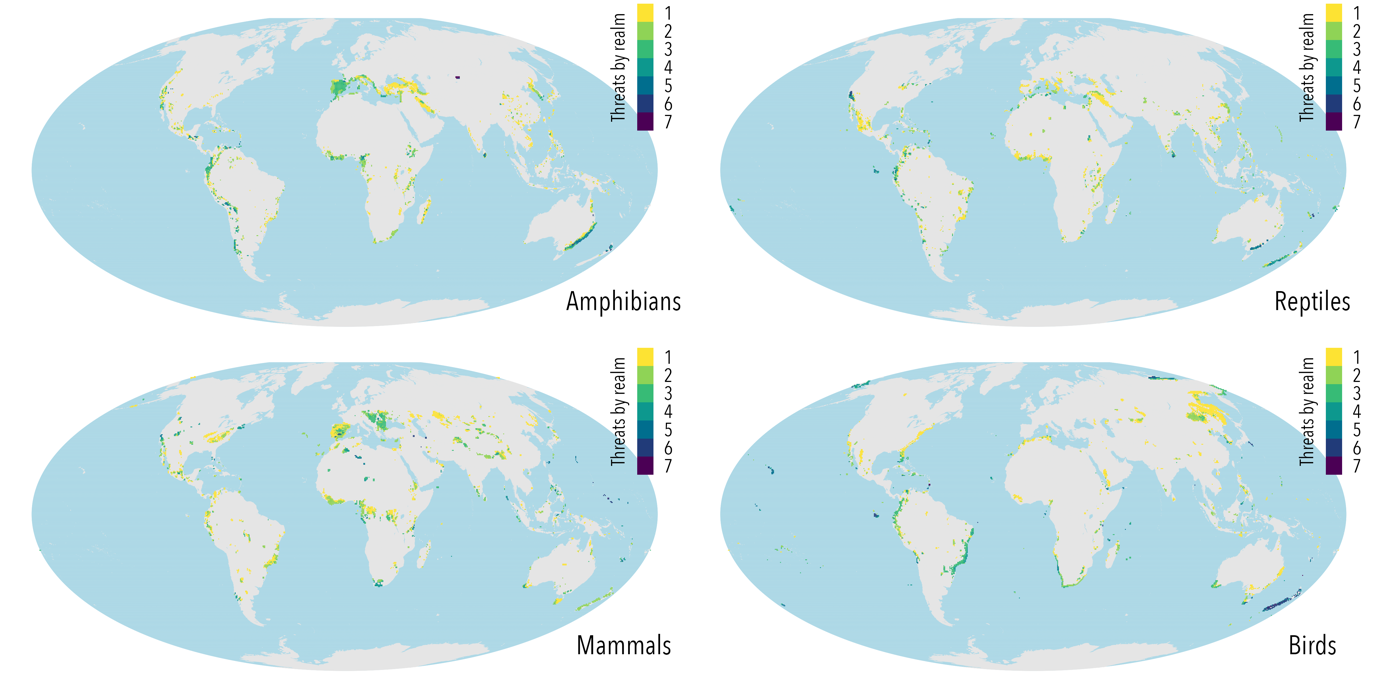

**Figure S8** | Number of hotspots at the regional level coloured by the number of threats that match the 10th decile.

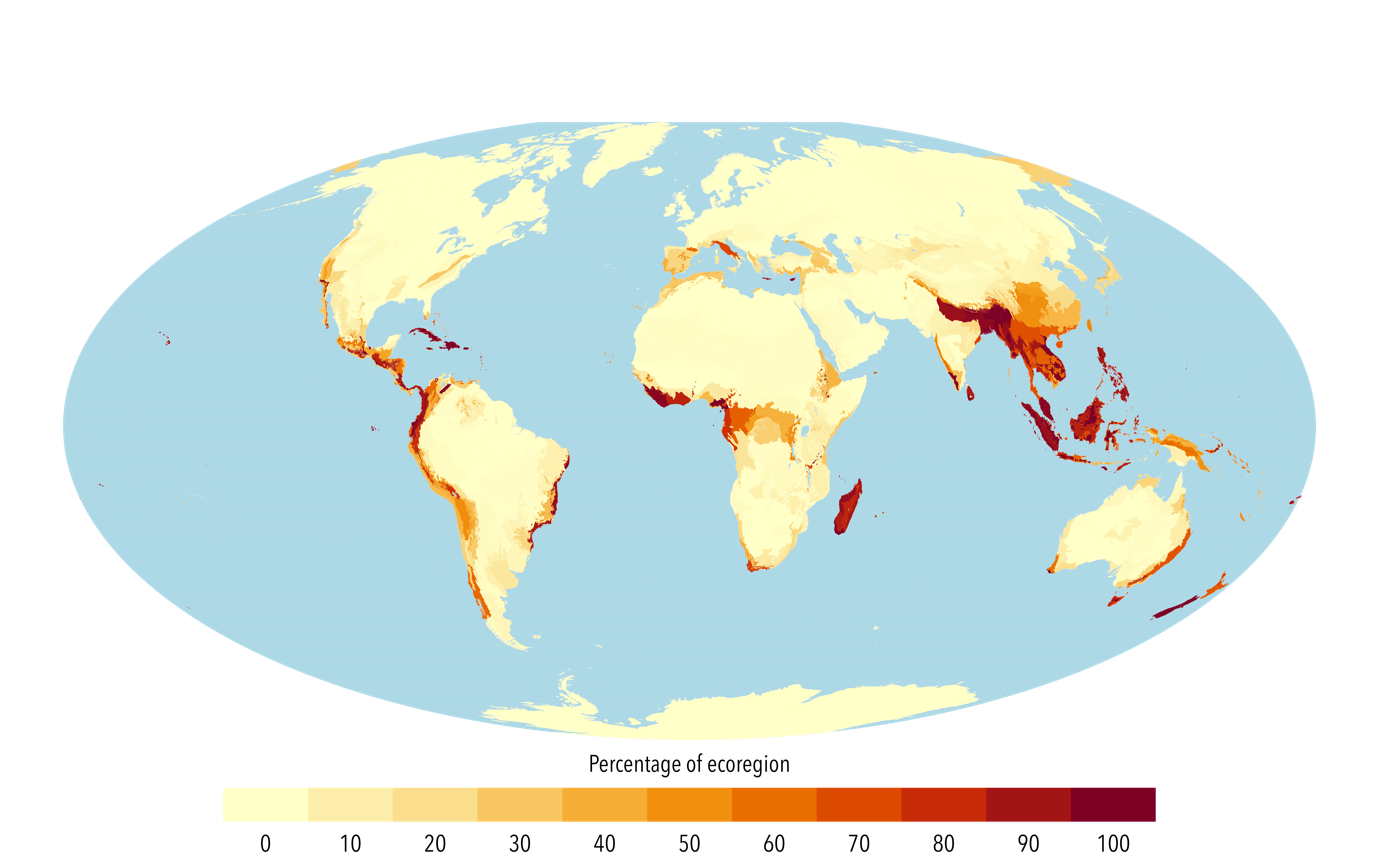

**Figure S9** | The percentage of the ecoregion that has hotspot cells, calculated at the pixel level. A value of 100 corresponds to the totality of the pixels being considered a hostpot.

**Table S1** | Number of grid cells in the top 10 quantile of the threats and WEGE for each by taxonomic groups.

| taxon | agriculture | logging | hunting | pollution | invasives | climate_change | urbanization |
| --- | --- | --- | --- | --- | --- | --- | --- |
| amphibians | 969 | 1378 | 334 | 180 | 1038 | 400 | 614 |
| reptiles | 244 | 2129 | 92 | 387 | 821 | 368 | 119 |
| mammals | 2533 | 3047 | 830 | 194 | 510 | 488 | 1381 |
| birds | 1359 | 2690 | 234 | 144 | 712 | 194 | 623 |

**Table S2** | Vertebrate hotspots across biomes.

| Biomes | People in biome | People in hotspot | Area in biome (KM^2^) | Area in hotspot (KM^2^) | Percentage of people in biome | Percentage of people globally | Percentage of land in biome | Percentage of hotspot in biome |
| --- | --- | --- | --- | --- | --- | --- | --- | --- |
| Boreal Forests/Taiga | 31,096,731 | 15,366 | 16,567,500 | 17,500 | 0% | 0% | 0% | 0.1% |
| Deserts & Xeric Shrublands | 806,653,953 | 86,652,895 | 27,860,000 | 1,192,500 | 10.7% | 1.1% | 0.7% | 4.3% |
| Flooded Grasslands & Savannas | 101,248,504 | 6,878,498 | 1,205,000 | 30,000 | 6.8% | 0.1% | 0% | 2.5% |
| Mangroves | 141,707,053 | 63,636,273 | 1,720,000 | 755,000 | 44.9% | 0.8% | 0.4% | 43.9% |
| Mediterranean Forests, Woodlands & Scrub | 358,317,811 | 151,624,758 | 4,525,000 | 1,090,000 | 42.3% | 1.9% | 0.6% | 24.1% |
| Montane Grasslands & Shrublands | 159,843,640 | 42,108,805 | 4,905,000 | 815,000 | 26.3% | 0.5% | 0.5% | 16.6% |
| Rock and Ice | 756,176 | 179,597 | 16,780,000 | 17,500 | 23.8% | 0% | 0% | 0.1% |
| Temperate Broadleaf & Mixed Forests | 1,780,555,080 | 250,939,129 | 15,042,500 | 1,882,500 | 14.1% | 3.2% | 1.1% | 12.5% |
| Temperate Conifer Forests | 148,429,785 | 17,370,132 | 4,175,000 | 290,000 | 11.7% | 0.2% | 0.2% | 6.9% |
| Temperate Grasslands, Savannas & Shrublands | 342,604,956 | 22,324,164 | 11,065,000 | 410,000 | 6.5% | 0.3% | 0.2% | 3.7% |
| Tropical & Subtropical Coniferous Forests | 104,098,807 | 72,043,993 | 740,000 | 392,500 | 69.2% | 0.9% | 0.2% | 53% |
| Tropical & Subtropical Dry Broadleaf Forests | 604,029,884 | 174,789,468 | 4,617,500 | 1,580,000 | 28.9% | 2.2% | 0.9% | 34.2% |
| Tropical & Subtropical Grasslands, Savannas & Shrublands | 927,379,233 | 78,592,868 | 22,212,500 | 895,000 | 8.5% | 1% | 0.5% | 4% |
| Tropical & Subtropical Moist Broadleaf Forests | 2,385,750,260 | 1,731,503,642 | 25,420,000 | 11,497,500 | 72. 6% | 21.9% | 6.8% | 45.2% |
| Tundra | 2,704,699 | 47,168 | 12,535,000 | 307,500 | 1.7% | 0% | 0.2% | 2.5% |

Dor each biome, the total estimated population and the one within the hotspots, the area total areas occupied by the biome and the area occupied by the hotspots and the percentages of people in hotspots, people at the global scale, the percentage of land and the percentage of hotspot area within each biome. The values were produced by agregating the 1kmx1km population count layer to a 50x50km grid to match the resolution used in this study.

**Table S3** | Vertebrate hotspots across the original biodiversity hotspots.

| CI HOTSPOTS | Hotspot pixel counts | Hotspot pixel counts total | Percentage of hotspot pixels in hotspost area |
| --- | --- | --- | --- |
| Sundaland | 836 | 952 | 87,8% |
| New Zealand | 179 | 204 | 87,7% |
| Madagascar and the Indian Ocean Islands | 304 | 356 | 85,4% |
| Guinean Forests of West Africa | 213 | 255 | 83,5% |
| Philippines | 265 | 331 | 80,1% |
| Caribbean Islands | 264 | 331 | 79,8% |
| Western Ghats and Sri Lanka | 59 | 74 | 79,7% |
| Tumbes-Choco-Magdalena | 124 | 159 | 78% |
| Indo-Burma | 884 | 1160 | 76,2% |
| Cape Floristic Region | 20 | 29 | 69% |
| Wallacea | 298 | 439 | 67,9% |
| Forests of East Australia | 67 | 100 | 67% |
| Polynesia-Micronesia | 243 | 427 | 56,9% |
| Himalaya | 166 | 304 | 54,6% |
| East Melanesian Islands | 129 | 237 | 54,4% |
| New Caledonia | 25 | 46 | 54,3% |
| Tropical Andes | 325 | 621 | 52,3% |
| Chilean Winter Rainfall and Valdivian Forests | 107 | 221 | 48,4% |
| Madrean Pine-Oak Woodlands | 70 | 174 | 40,2% |
| Mesoamerica | 241 | 611 | 39,4% |
| California Floristic Province | 57 | 150 | 38% |
| Succulent Karoo | 15 | 42 | 35,7% |
| Mountains of Southwest China | 38 | 108 | 35,2% |
| Atlantic Forest | 168 | 491 | 34,2% |
| Eastern Afromontane | 126 | 396 | 31,8% |
| Japan | 89 | 337 | 26,4% |
| Coastal Forests of Eastern Africa | 29 | 117 | 24,8% |
| Mediterranean Basin | 303 | 1242 | 24,4% |
| Southwest Australia | 30 | 143 | 21% |
| Maputaland-Pondoland-Albany | 21 | 110 | 19,1% |
| Caucasus | 28 | 209 | 13,4% |
| Mountains of Central Asia | 38 | 348 | 10,9% |
| Irano-Anatolian | 29 | 364 | 8% |
| Horn of Africa | 51 | 784 | 6,5% |
| NA | 2,604 | 54,568 | 4,8% |
| North American Coastal Plain | 13 | 490 | 2,7% |
| Cerrado | 11 | 817 | 1,3% |

The number and percentage of pixels of the hotspot pixels generated in tihis study by the CI hotspots converted to pixel. A high percentage represents a high proportion of pixels of the CI hotspots that were also considered hotspots in this analysis.

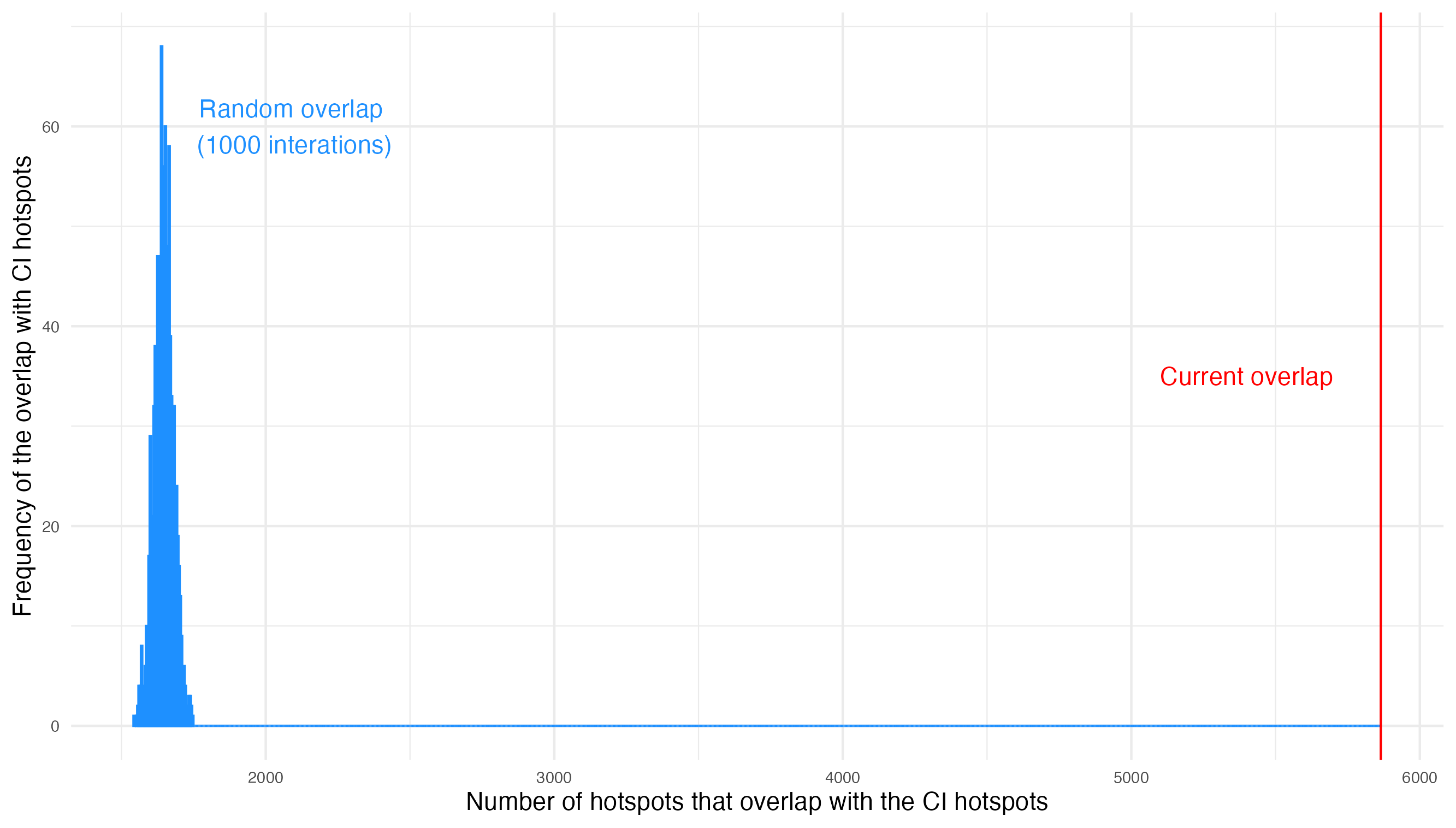

**Figure S10** | A figure showcasing the random overlap of terrestrial cells with the CI hotspots in blue with a mean of 1645 grids (19%) and the current overlap of our generated hotspots in red with a value of 5865 grids (64% of all grids).
